# Environmental Regulation and Gene-by-Environment Interaction Influence *RAP1* Activity and its Impact on Gene Expression

**DOI:** 10.64898/2026.05.06.723246

**Authors:** Siddhant Kalra, Guadalupe Sanchez, Alexandra Stubin, Anya Le, Arpie Bakshian, Bianca Ortiz Diaz, Brianna Meiko Mark, Crystal Peña, Eliza Parker, Emma Johnston, Evan Edward Hsu, Gavin Brangham, Idenya Bala-Mehta, Luis Daniel Perez, Maya Lee Milrod, Meili Stanten, Mikoto Nakamura, Pete Hwang, Sara Maria Ptaszynska, Sasha Cander, Sebin Park, Tai Lon Tan, Yancheng Zhou, Joseph David Coolon

## Abstract

Gene-by-environment (GxE) interactions play a major role in shaping both phenotypic and molecular variation, with important implications for human health and disease. In this study, we used the Doxycycline (Dox) regulated, tetracycline-responsive (Tet-Off) promoter system to sequentially reduce or titrate gene expression levels of the essential yeast transcription factor *Repressor Activator Protein 1* (*RAP1*) similar to a hypomorph allele series, across three distinct environments: Yeast Peptone Dextrose (YPD) media, YPD media with Heat Shock (HS), and Yeast Peptone Acetate (YPAC) media. We then performed RNA sequencing (RNA Seq) to assess global transcriptional responses to *RAP1* reduction in these different growth environments. Our analysis first focused on the independent effects of varying *RAP1* expression levels within and across environments. We then explored GxE interactions, revealing a subset of genes with significant consequences of reduced levels of *RAP1* and environment-specific expression patterns. Notably, many genes exhibited opposite effects of *RAP1* titration on gene expression when yeast were grown in YPAC media compared to YPD media and/or HS, suggesting environment-dependent regulatory architecture. This design reveals how cells integrate internal transcriptional and regulatory changes with external environmental cues, providing a deeper view of GxE architecture. Using Weighted Gene Co-expression Network Analysis (WGCNA), we identified co-regulated gene modules, and by combining this with transcription factor motif enrichment tests, our study identified candidate regulators driving their dynamics. Our findings demonstrate that gene regulatory networks can vary dramatically depending on the environmental context an organism experiences, which can then influence the specific phenotypes produced by a particular genetic perturbation. This illustrates the complexity of genotype–environment interactions and the importance of studying gene function in multiple environments to gain a truly comprehensive understanding of a gene’s sometimes numerous and diverse functions.

## Introduction

Living organisms must maintain internal homeostasis despite exposure to dynamic environmental conditions. Sensing and responding to external stimuli are essential for survival and evolutionary fitness^1^. At the molecular level, cells achieve this plasticity through complex and intricate signal transduction networks that detect environmental cues and initiate cascades of regulatory events, that typically modulate gene-specific gene expression levels^2,3^. These regulatory mechanisms not only determine transcriptional activation and repression but also fine-tune the magnitude of gene expression at transcriptional, post-transcriptional, translational, and epigenetic levels, ultimately contributing to phenotypes and phenotypic diversity^4–9^. This complex regulatory architecture confers flexibility, enabling organisms to thrive under favorable conditions but also respond to environmental stresses like extreme temperature and altered nutrient availability. This phenomenon known as Gene-by-Environment (GXE) interaction has been recognized for decades^10–14^ to form a complex interplay between an individual’s genetic makeup and the environment encountered both prenatally and throughout life^15,16^. These mechanisms have evolved not only to ensure immediate survival but also to foster long-term resilience and evolutionary fitness across generations^17^.

Whether a gene is expressed and the extent to which it is expressed are strongly shaped by the environment an organism experiences, and this has important implications for both trait expression and disease states^18^. A classic example of GXE interaction is the genetic disorder Phenylketonuria (PKU), which is caused by mutations in the *PAH* gene, leading to a buildup of phenylalanine^19^. When individuals with PKU consume a high-phenylalanine diet, it can cause severe cognitive damage, but early dietary restriction can significantly reduce these effects^20^. This example not only highlights the significance of GXE interactions but also points out that such interactions are often sensitive during a particular developmental window.

The use of high-throughput transcriptomics technologies such as RNA seq has enabled the study of dynamic gene expression patterns across species and ecosystems. As personal genomics becomes increasingly incorporated in healthcare, understanding how environmental factors modulate gene expression is of critical importance. This knowledge not only informs predictions about individual health trajectories but also supports the design of targeted interventions, such as environmental modifications, dietary adjustments, or gene-targeted therapies. Furthermore, examining these GXE interactions within an ecological context advances our understanding of how organisms adapt, survive, and evolve in changing environments.

*Saccharomyces cerevisiae* is a powerful system to study GXE interactions due to its well-characterized genome, rapid growth rate, and the ease of both genetic manipulation and environmental control. Pioneering studies by Brem et al.^21^ and Smith et al.^22^ utilized quantitative trait locus (QTL) mapping in yeast to identify genetic loci underlying phenotypic variation across multiple environmental contexts. Notably, the phenotypic consequences of these alleles were shown to be environment-dependent, illustrating the conditional nature of genetic effects. Collectively, these findings reinforce the concept that complex traits are shaped by a dynamic interplay between genetic background and environmental conditions.

Yeast Extract, Peptone, Dextrose (YPD) is a nutrient-rich growth medium commonly used for culturing *S. cerevisiae*. It contains glucose (as dextrose) as the primary carbon source that supports rapid cell proliferation, with typical doubling times of approximately 90 minutes during exponential growth under optimal conditions. This rapid growth reflects the highly efficient fermentative metabolism of *S. cerevisiae*, which preferentially utilizes glucose through glycolysis, even in the presence of oxygen (a phenomenon known as the Crabtree effect)^23^. However, environmental perturbations such as elevated temperature can severely compromise cellular functions. Heat shock can lead to protein misfolding and aggregation, disrupt cellular structures, and potentially cause DNA damage, thereby compromising cell viability^24^. To combat these stressors, *S. cerevisiae* initiates a conserved transcriptional program known as the Heat Shock Response (HSR), primarily mediated by heat shock transcription factors like Hsf1^25,26^. In contrast to glucose-rich conditions, *S. cerevisiae* is also capable of adapting to growth on alternative carbon sources such as acetate. This metabolic switch is particularly relevant in nutrient-limited conditions, where glucose levels can become depleted^27^. Growth on acetate necessitates a fundamental reorganization of the metabolic landscape in yeast cells. Glycolytic genes are transcriptionally repressed, while genes involved in the tricarboxylic acid (TCA) cycle, the glyoxylate shunt, gluconeogenesis, and oxidative phosphorylation are upregulated to support energy production through respiration^27,28^. Together, the ability of *S. cerevisiae* to grow under diverse nutritional and stress conditions, ranging from fermentative growth in YPD, to stress adaptation during heat shock, to respiratory metabolism on acetate, makes it an ideal model system for studying gene-environment interactions and regulatory plasticity.

Traditional gene expression studies typically examine how transcriptomic profiles shift across environmental conditions while maintaining a constant genetic background^29,30^, or they compare different genetic backgrounds within the same environment^21,31^. Such approaches capture only a subset of possible dynamics and frequently miss gene–environment (GxE) interactions. Far fewer studies systematically vary both genetic makeup and environment conditions simultaneously^32–34^. In this work, we expand on this framework by introducing an additional regulatory dimension to dissect GXE interactions by systematically modulating the levels of a key transcriptional regulator, Repressor Activator Protein 1 (*RAP1*), using a Doxycycline (Dox) regulated Tet-off system. Rap1p is an essential transcription factor in *S. cerevisiae*, known for its dual role as an activator and/or repressor depending on genomic context and associated cofactors^35^. Rap1p binds to ∼5% of yeast genome where its activity is essential for silencing, metabolic regulation, and stress responses through interactions with Sir proteins^35–38^ and other regulatory elements along with its recently discovered role in splicing^39^.

In this study we performed RNA seq on a yeast strain in which the endogenous *RAP1* promoter was replaced with a Tet-Off promoter system (Figure 1A), enabling precise titration of *RAP1* levels depending on the concentration of Dox in their media such that increasing concentrations of Dox resulted in progressively lower levels of *RAP1* expression^35,40^ (Figure 1B). This systematic series of yeast cultures with varied levels of reduction in *RAP1* expression levels behaves quite similar to a hypomorph allele series, and we use this here to represent different genetic states. By titrating *RAP1* expression level with multiple Dox concentrations, we introduce controlled variation in *RAP1* transcriptional output, which we intersect with three ecologically and metabolically distinct environments: rich media (YPD), heat shock (HS), and media with an alternative carbon source (YPAC) (Figure 1 C, D) (S-Table 1). This experimental design allows interrogation of cellular integration of internal transcriptional regulatory perturbation with external environmental conditions, offering a nuanced understanding of GXE architecture. Furthermore, we employed weighted gene co-expression network analysis (WGCNA), enabling the identification of co-regulated genes resulting in different network modules^41^. Moreover, by integrating WGCNA and transcription factor (TF) motif enrichment analysis, we identified candidate TFs likely orchestrating the dynamic behavior of these modules, further refining our understanding of the regulatory logic underlying GXE responses. Together, these findings offer a systems-level perspective of how transcriptional networks dynamically change in response to the combined effects of genetic and environmental perturbations, uncovering context-specific regulatory mechanisms that shape phenotypic outcomes.

**Figure 1:**
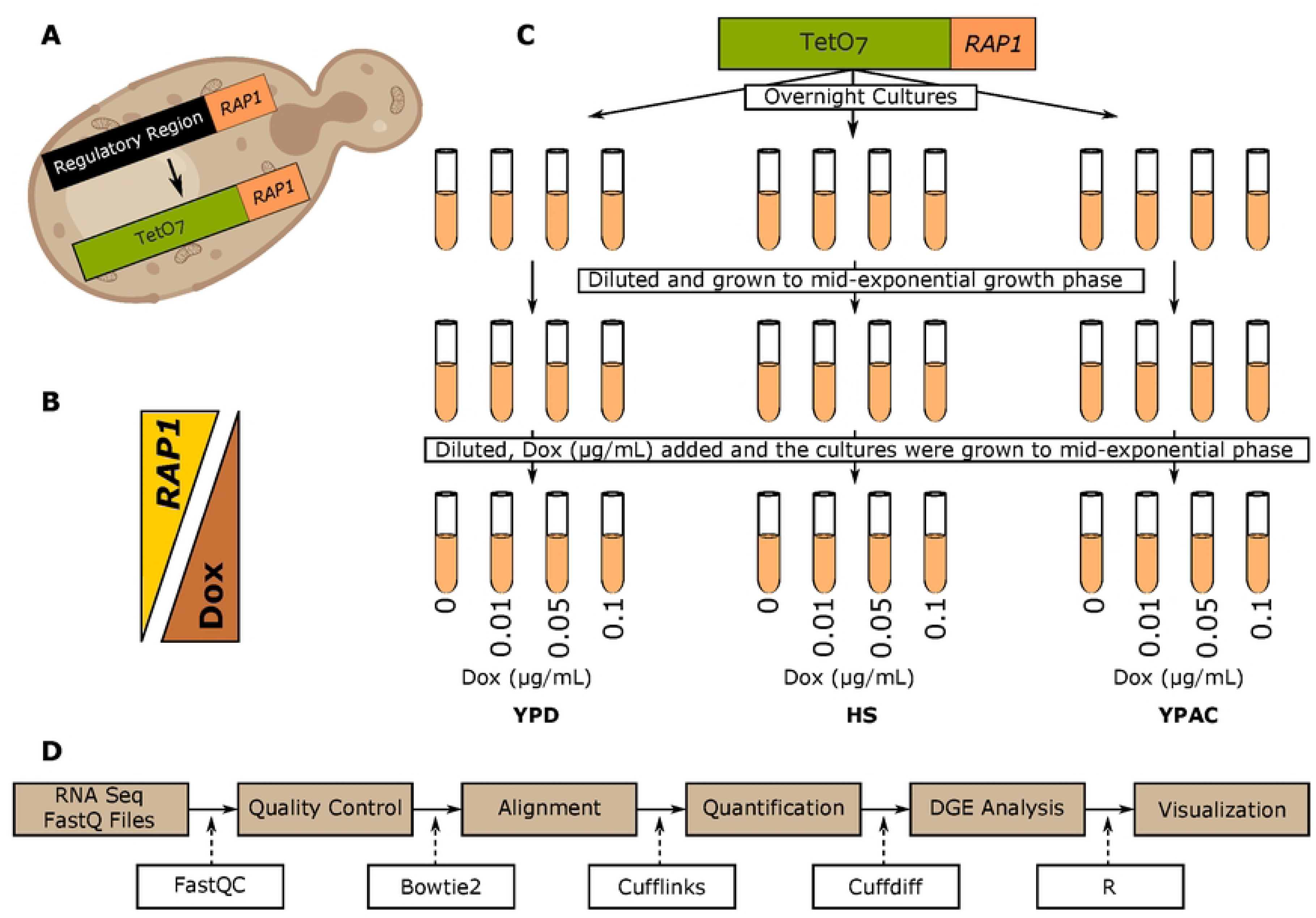
Experimental design and analysis workflow. A) Overview of the Tet-Off system used to modulate *RAP1* levels. B) Inverse relationship between Dox concentration in the media and *RAP1* expression levels. C) Culture scheme showing *RAP1* titration across multiple Dox levels in three growth environments. D) Computational pipeline used for downstream data processing.

## Results

### Environment-driven gene expression differences

To characterize environment-specific transcriptional responses of yeast with unmodified *RAP1* expression levels (zero Dox), we performed differential gene expression analyses of yeast grown in YPD, HS, and YPAC conditions (Figure 2 A-F). The HS vs YPD comparison identified 806 significantly differentially expressed genes (DEGs) (R^2^ = 0.98, *P* < 0.001), including 655 upregulated (log_2_ fold change > 0) and 151 downregulated genes (log_2_ fold change < 0) (Figure 2A, D). The YPAC vs YPD comparison found a substantially broader response, with 3727 DEGs (2070 upregulated and 1657 downregulated) (R^2^ = 0.19, *P* < 0.001) (Figure 2B, E). Finally, comparing YPAC to HS revealed 3743 DEGs (R^2^ = 0.23, *P* < 0.001), comprising 1899 upregulated and 1844 downregulated transcripts (Figure 2C, F). These contrasts highlight the distinct transcriptional programs engaged by each environment at unmodified *RAP1* levels.

**Figure 2.**
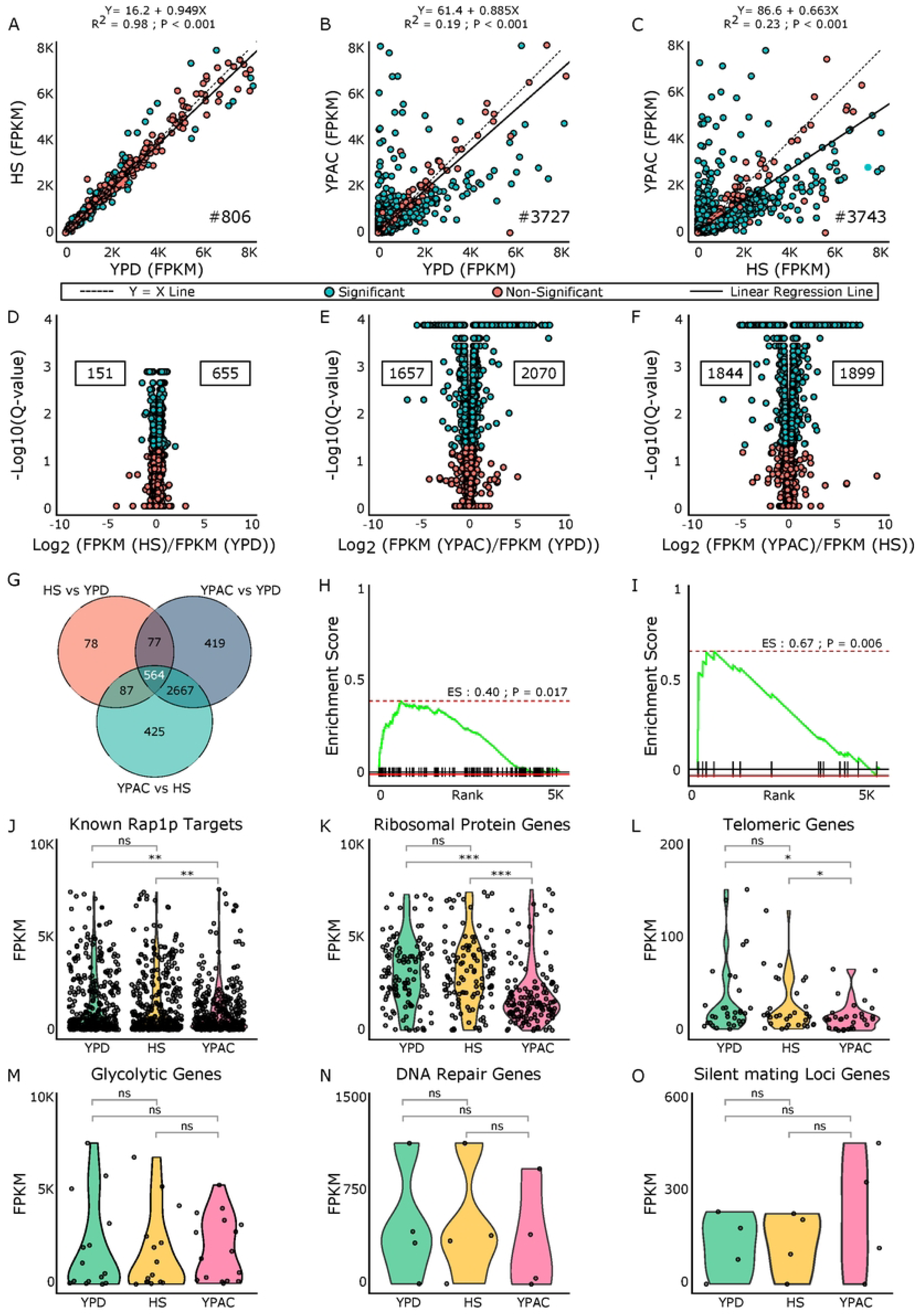
Differential expression patterns at maximal *RAP1* levels across environments. Scatter plots comparing gene expression at max *RAP1* levels across conditions: A) HS vs YPD, B) YPAC vs YPD, and C) YPAC vs HS. Blue points represent genes with significant changes while red points denote nonsignificant genes. The black dashed line shows Y=X line, and the solid black line shows the line of best fit from linear regression. Each panel reports the regression equation, R², and p value on the top along with the count of significant DEGs noted in the lower right. D–F) Volcano plots showing up and downregulated genes for each comparison. Values on the right indicate the number of significantly upregulated genes, and values on the left indicate significantly downregulated genes. G) Overlap of significant DEGs across the three comparisons. Gene set enrichment analysis for known H) heat-shock and I) acetate-responsive gene sets, evaluated against DEGs from HS vs YPD and YPAC vs YPD, respectively. Gene expression profile comparison for J) *RAP1* targets, K) ribosomal protein genes, L) telomeric genes, M) glycolytic genes, N) DNA repair genes, and O) silent mating–locus genes across YPD, HS, and YPAC.

Among the identified differentially expressed genes, we found a core set of 564 genes shared across all three environmental comparisons, indicating a common transcriptional response to these environmental changes (Figure 2G) (S-Table2). Beyond this shared set, 78 genes responded exclusively to the shift from YPD to HS, while 419 genes were uniquely affected by growth in media containing different carbon sources (YPD vs YPAC). An additional 425 genes were specific to the HS vs YPAC comparison (Figure 2G) (S-Table2). The DEGs from HS and YPAC showed clear overlap with genes linked to these conditions in literature (see methods) (S-Table3). The DEGs responding to HS displayed moderate enrichment (Enrichment Score (ES) = 0.40, *P* = 0.017), with known HS responding genes (Figure 2H). In contrast, the DEGs responding to growth in YPAC media showed stronger enrichment for genes involved in metabolism (ES = 0.67, *P* = 0.006) (Figure 2I), suggesting that the transcriptional changes we observe under respiratory growth (YPAC) more closely match established acetate associated signatures.

For our lists of significant DEGs responding to different environmental conditions, we also performed gene ontology (GO) term enrichment analysis (S-Table 4) and examined the overlap of enriched terms (S-Figure 1). The 16 enriched GO terms exclusive to the comparison of HS vs YPD conditions were primarily associated with iron transport and storage including transition metal ion transport, vacuolar storage, intracellular iron homeostasis, and iron coordination entity transport (S-Figure 1) (S-Table 4). The 64 terms enriched exclusively in comparison of YPAC to YPD growth conditions were strongly associated with respiration and mitochondrial biogenesis, such as the mitochondrial outer membrane, energy-coupled proton transmembrane transport, and oxidoreductase activity (S-Figure 1) (S-Table4). The 137 terms enriched exclusively in YPAC relative to HS growth conditions encompassed diverse biological processes. These included pathways associated with transcriptional regulation and growth, such as the RNA polymerase I transcription regulator complex and DNA-binding transcription factor activity, alongside metabolic, developmental, and biosynthetic processes, including long-chain fatty acid transport, ascospore formation, developmental processes, and ribosome biogenesis. (S-Figure 1) (S-Table 4).

Next, we examined how the known regulatory functions of *RAP1* are affected by growth in different environments (Figure 2J-O). Using gene subsets (known downstream targets of *RAP1*, Ribosomal protein genes (RPGs), Telomeric genes, Glycolytic genes, DNA repair genes, and Silent mating loci (SML)) defined in our previous study^35^, we compared their expression profiles (FPKM values) across the three conditions (YPD, HS, and YPAC) at unmodified *RAP1* levels. This analysis allowed assessment of whether *RAP1’s* known regulatory roles are maintained, enhanced, or altered in response to different environmental contexts. We observed a significant change in the expression of known *RAP1* downstream targets in YPAC compared to both YPD and HS conditions. In contrast, their expression remained largely unchanged in HS relative to YPD (Figure 2J). This trend was consistent for the ribosomal protein gene (RPG) family (Figure 2K) and telomeric genes (Figure 2L). Specifically, we observed a significant change in the expression of RPGs and telomeric genes in YPAC compared to both YPD and HS conditions, but their expression remained largely unchanged in HS relative to YPD. We didn’t observe any significant change in gene expression for glycolytic (Figure 2M), DNA repair (Figure 2N), and silent mating loci (SML) genes in any of our environmental comparisons (Figure 2O).

### Identifying gene expression responses to *RAP1* titration in different growth environments

Analysis of genome-wide gene expression in yeast with unmodified *RAP1* expression (0 Dox) grown in different environmental conditions identified numerous genes with significant changes in expression level. Next, we sought to identify those genes that had significant GxE interactions. To do this we used the Tet-Off system to sequentially reduce *RAP1* expression levels by increasing Dox concentrations in the growth media and assessed genome-wide gene expression changes with RNA seq (Figure 1 A-D) (S-Table1). In the YPD environment (Figure 3A-F), reducing *RAP1* levels with 0.01 µg/mL Dox (compared to the control with no Dox) resulted in 756 significantly DEGs (R² = 0.98, p < 0.001) (Figure 3A), including 239 downregulated (log2 fold change < 0) and 517 upregulated genes (log2 fold change > 0) (Figure 3D). When *RAP1* levels were further reduced (with 0.05 µg/mL Dox), we identified 1338 significant DEGs (R² = 0.95, p < 0.001) (Figure 3B), with 478 downregulated and 860 upregulated genes identified (Figure 3E). Upon the greatest *RAP1* depletion tested (0.1 µg/mL Dox), the number of significant DEGs increased to 1748 (Figure 3C), where 740 genes were downregulated and 1008 were upregulated (Figure 3F). Overall, in the YPD environment, we observed that sequentially reducing *RAP1* expression levels led to an increasing number of significant DEGs (Figure 3A-F), which is consistent with findings from our previous study^35^.

**Figure 3.**
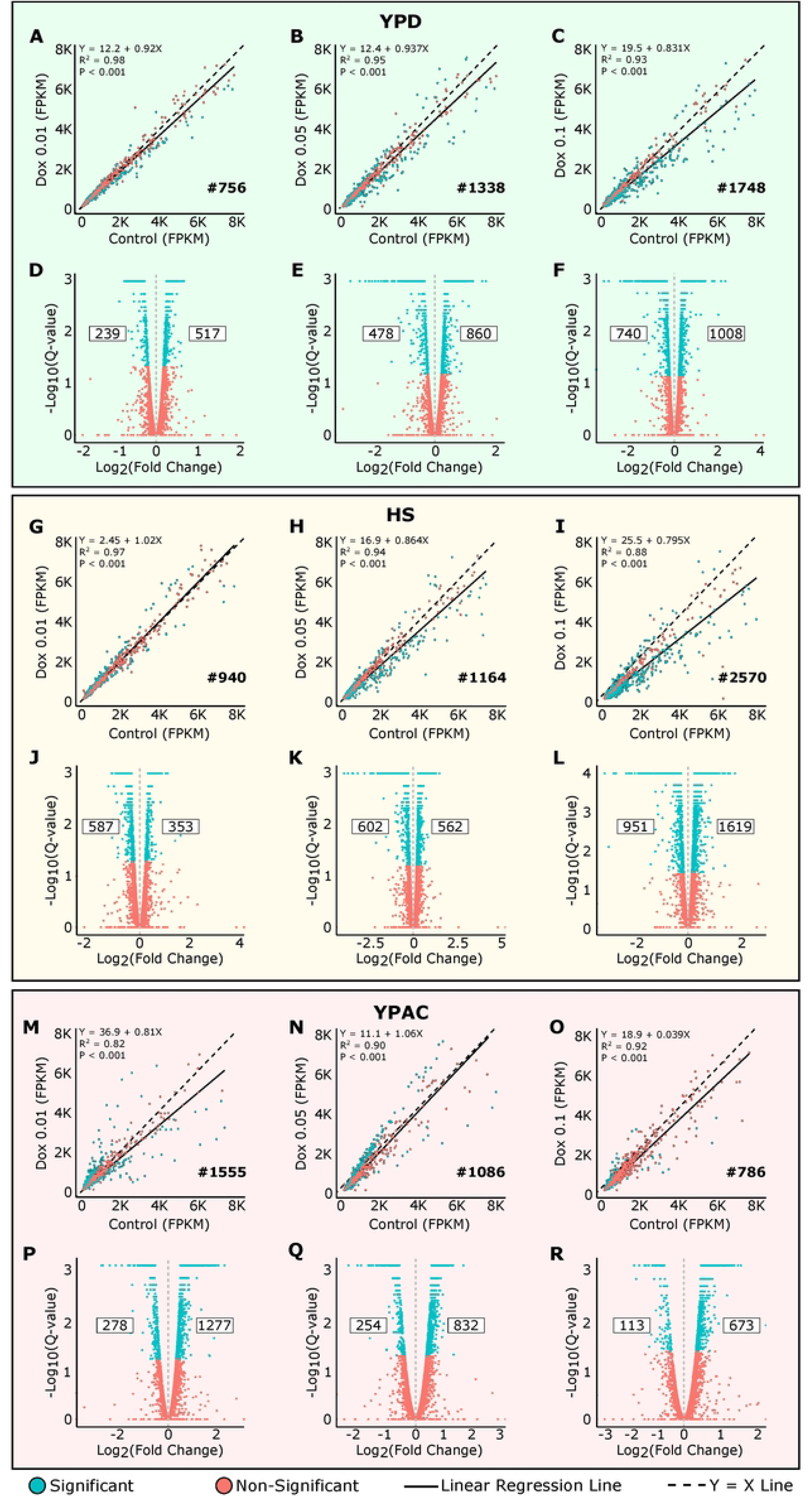
Differential expression patterns at multiple *RAP1* levels across environments. A-C) Scatter plots comparing gene expression at different *RAP1* levels compared to control in YPD environment. D–F) Volcano plots showing up and downregulated genes for each comparison in YPD. G-I) Scatter plots comparing gene expression at different *RAP1* levels compared to control in HS environment. J-L) Volcano plots showing up and downregulated genes for each comparison in HS. M-O) Scatter plots comparing gene expression at different *RAP1* levels compared to control in YPAC environment. P-R) Volcano plots showing up and downregulated genes for each comparison in YPAC. Blue points represent genes with significant changes while red points denote nonsignificant genes. In each scatter plot black dashed line shows Y=X line, and the solid black line shows the line of best fit from linear regression. Each panel reports the regression equation, R², and p value on the top along with the count of significant DEGs noted in the lower right. For each volcano plot values on the right indicate the number of significantly upregulated genes, and values on the left indicate significantly downregulated genes.

When yeast was given HS treatment (Figure 3G-L), reducing *RAP1* levels with 0.01 µg/mL Dox resulted in 940 significantly DEGs (R² = 0.97, p < 0.001) (Figure 3G), including 587 downregulated and 353 upregulated genes identified (Figure 3J). When *RAP1* levels were further reduced with 0.05 µg/mL Dox, we identified 1164 significant DEGs (R² = 0.94, p < 0.001) (Figure 3H), with 602 downregulated and 562 upregulated genes found (Figure 3K). Upon the highest *RAP1* depletion (0.1 µg/mL Dox), the number of significant DEGs increased to 2570 (R² = 0.88, p < 0.001) (Figure 3I), of which 951 were downregulated and 1619 were upregulated (Figure 3L). The HS environment, similar to that observed for YPD, resulted in an overall increase in number of DEGs as *RAP1* levels were sequentially reduced.

Growth in YPAC media (Figure 3M-R) while modulating *RAP1* levels using the Tet-Off system revealed distinct gene expression changes that were opposite to those observed in YPD and HS. Reducing *RAP1* levels with 0.01 µg/mL Dox, resulted in 1555 significant DEGs (R² = 0.82, p < 0.001) (Figure 3M), with 278 downregulated and 1277 upregulated genes identified (Figure 3P). Further reduction of *RAP1* levels with 0.05 µg/mL Dox led to 1086 significant DEGs (R² = 0.90, p < 0.001) (Figure 3N), comprising 254 downregulated and 832 upregulated genes (Figure 3Q). At the lowest *RAP1* level (0.1 µg/mL Dox), the number of significant DEGs decreased to 786 (R² = 0.92, p < 0.001) (Figure 3O), with 113 found to be downregulated and 673 upregulated (Figure 3R). Overall, in the YPAC condition, we observed that progressively reducing *RAP1* levels led to a decreasing number of significant DEGs observed. These findings indicate that the impact of *RAP1* depletion on gene expression is strongly influenced by the environmental context.

### Comparative analysis of transcriptional responses to altered *RAP1* levels across environments

In the YPD environment, we observed distinct gene expression responses to varying levels of *RAP1* (Figure 4A). Specifically, 617 genes responded exclusively to Dox 0.1; 269 genes were uniquely responsive to Dox 0.05, and 118 genes responded only to Dox 0.01. When identified DEG overlap across Dox levels was determined, we found 546 genes were shared between the Dox 0.1 and Dox 0.05 conditions, while 115 genes were common to both Dox 0.1 and Dox 0.01. A smaller subset of 53 genes overlapped between the Dox 0.05 and Dox 0.01 conditions. Notably, 470 genes exhibited significant differential expression across all three Dox concentrations, suggesting a core set of genes that are consistently responsive to *RAP1* perturbation regardless of dosage (Figure 4A).

**Figure 4.**
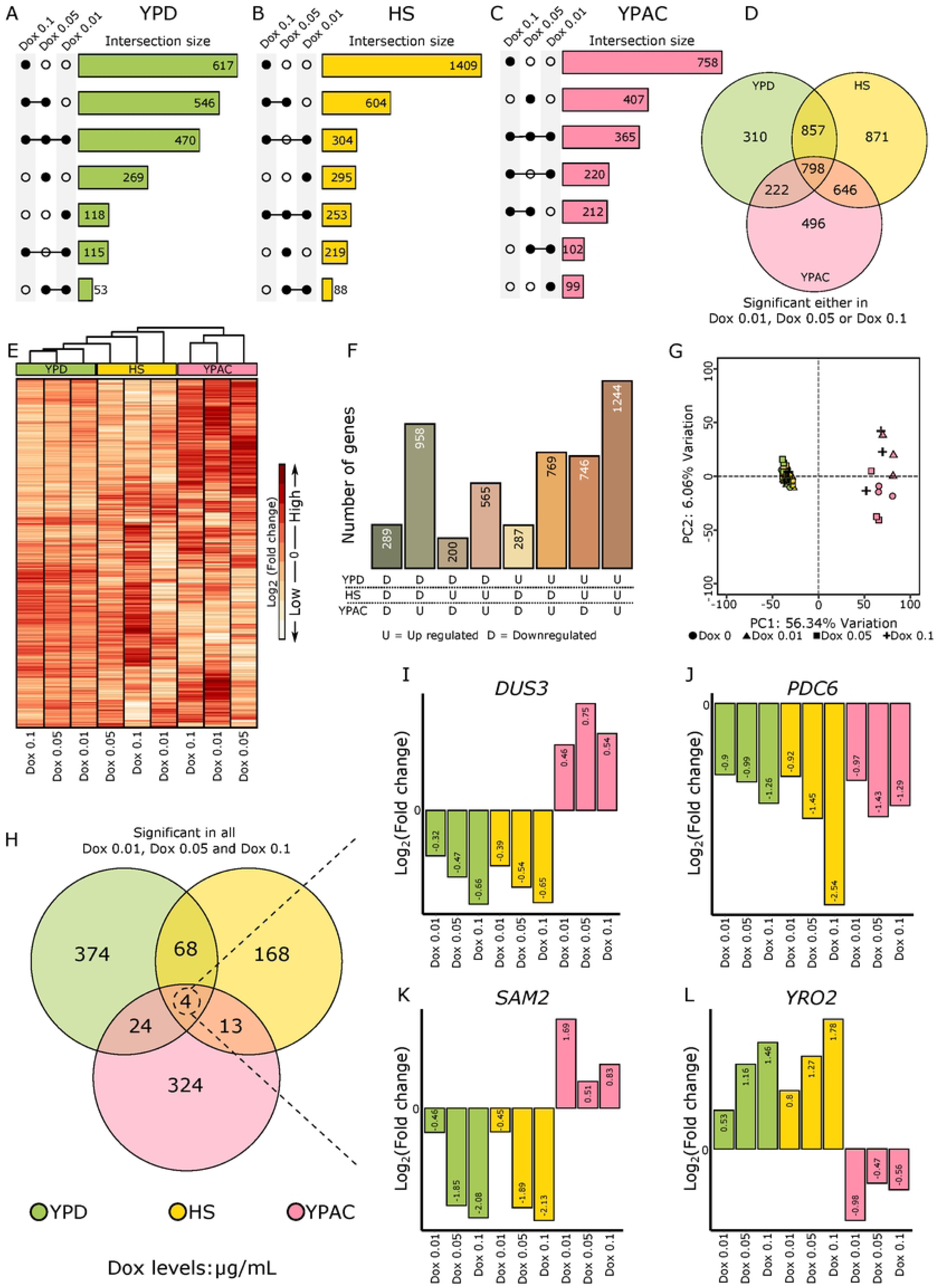
Gene by environment effects. Upset plots showing the overlap of significant DEGs across *RAP1* levels within each environment: A) YPD, B) HS, and C) YPAC. D) Venn diagram summarizing genes that were significant at any *RAP1* level across the three environments. E) Heatmap of log_2_ fold change values illustrating how *RAP1*-dependent responses vary across YPD, HS, and YPAC. F) Quantification of all possible direction of change combinations across the three environments. G) PCA plot showing variation explained by the first two principal components. H) Overlap of significant DEGs shared across all RAP1 levels in the three environments. I–L) Bar plots showing log_2_ fold change values for four genes that remained significant across all *RAP1* levels in all environments.

Within the HS environmental condition, exposure to Dox 0.1 resulted in 1409 DEGs exclusive to this Dox concentration, reflecting a strong transcriptional response at the lowest *RAP1* levels. In comparison, 295 genes responded solely at Dox 0.01, and 219 genes were uniquely responsive to Dox 0.05. When examining the overlap between conditions, 604 genes were shared between Dox 0.1 and Dox 0.05; 304 genes shared between Dox 0.1 and Dox 0.01, and 88 genes were common to Dox 0.05 and Dox 0.01. 253 genes showed significant expression changes across all three Dox (*RAP1*) levels. This suggests that, under heat stress, the majority of *RAP1* responsive genes are activated at the lowest *RAP1* expression level (at Dox 0.1, Figure 4B).

In the YPAC environment, the highest Dox concentration (Dox 0.1) triggered exclusive differential expression of 758 genes, while 407 and 99 genes responded uniquely to the intermediate (Dox 0.05) and lowest (Dox 0.01) Dox concentrations respectively. 212 genes were shared between Dox 0.1 and Dox 0.05, 220 between Dox 0.1 and Dox 0.01, and 102 between Dox 0.05 and Dox 0.01. A total of 365 genes were consistently differentially expressed across all three Dox concentrations, highlighting a moderately sized core *RAP1* responsive gene set in this condition (Figure 4C).

Next, to investigate the extent of overlap in *RAP1* responsive genes across different environments, we examined how many genes were shared among YPD, HS, and YPAC conditions (Figure 4D). For this analysis, a gene was considered significant within an environment if it had significant differential expression at any Dox concentration/*RAP1* level. Using this criterion, we identified 798 genes that were common across all three environments, representing a core set of *RAP1* responsive genes. Pairwise comparisons revealed that 857 genes were shared between YPD and HS, 646 between HS and YPAC, and 222 between YPD and YPAC environments. Additionally, a substantial number of genes were uniquely responsive to *RAP1* levels in individual environments: 310 were exclusive to YPD, 871 exclusives to HS, and 496 exclusives to YPAC (Figure 4D). These findings highlight both conserved and environment-specific aspects of the transcriptional response to *RAP1* depletion, with HS showing the largest number of unique responsive genes.

We analyzed enriched GO terms among significant DEGs in each environmental condition independent of *RAP1* dosage and determined their overlap (S-Table5) (S-Figure 2). Across all three conditions, 44 GO terms were shared, while 82 terms were unique to growth in YPD, 41 were unique to HS, and 231 were unique to YPAC (S-Figure 2). The YPD exclusive GO terms included sulfate assimilation, carbon catabolite regulation, and ubiquitin ligase complex activity (S-Table5). These processes have been previously implicated in nutrient and growth-related responses in *Saccharomyces cerevisiae*, where abundant nutrients activate global nutrient-sensing and catabolite repression pathways that coordinate carbon and nitrogen metabolism and proteome allocation for growth and proliferation^42,43^. HS exclusive GO terms were dominated by rRNA processing and ribosome assembly (S-Table5). Ribosome biogenesis and associated rRNA processing underlie translational capacity and are transcriptionally regulated in response to growth and stress signals in *S. cerevisiae*^44^. The YPAC exclusive terms included mitochondrial and organellar ribosome functions (S-Table5). Mitochondrial and organellar ribosome functions reflect increased translation of components needed for oxidative phosphorylation, a key feature of respiratory metabolism in yeast when cells rely on mitochondrial energy production rather than fermentation^45^.

### Directional Effects of YPD and YPAC on Gene Expression

To get a broader view of gene expression response to *RAP1* depletion across the genome in different environments we visualized expression plasticity in the form of heatmap (Figure 4E) The YPAC elicits gene expression patterns often opposite to those observed in YPD and HS. The heatmap with hierarchical clustering based on log_2_FC values showed that YPD and HS share a similar expression pattern, while YPAC behaves quite differently. Most genes (1704) had an opposite direction of expression change in YPAC compared with both YPD and HS environments (Figure 4F). In fact, 958 genes that were downregulated as *RAP1* levels changed in both YPD and HS became upregulated in YPAC, and 746 genes that were upregulated in both YPD and HS became downregulated in YPAC highlighting the change in regulatory mode with the change in carbon source.

To evaluate overall transcriptional variation across conditions using a different methodology, we used Principal Component Analysis (PCA) and found it reinforced the conclusions from the hierarchical clustering (Figure 4G). PC1, which explained 56.43% of the variance, separates samples by environment. YPD and HS group tightly, whereas YPAC separates from both, pointing to the environment as the strongest determinant of transcriptional variation (Figure 4G). *RAP1* levels still play a significant regulatory role, determining which genes respond, along with the magnitude of response but the impact of *RAP1* is dependent on the environmental context. *RAP1’s* effects in YPD and HS move gene expression along a similar path, yet in YPAC, that same dosage change flips the response for many genes.

Next, we sought to identify genes that were consistently responsive across all environments and all Dox concentrations (Figure 4H). Applying this stringent criterion, we identified 374 genes exclusive to YPD, 168 exclusives to HS, and 324 exclusives to YPAC. Only four genes (*DUS3, PDC6, SAM2, and YRO2)* were significantly differentially expressed at all Dox levels across all three environments (Figure 4H). Visualizing their log₂ fold-change through bar plots (Figure 4I-L) revealed notable variation, not only in the magnitude of expression changes within the environment but also in the direction of the response, reflecting the combined influence of *RAP1* dosage and environmental context. Three out of these four genes (Figure 4I-L) exhibited opposite-direction responses across environments, again highlighting an environmental influence on *RAP1’s* dual role where it is acting as an activator for a gene in one environment and as a repressor for the same gene in different environment.

### Co-expression network analysis reveals modules associated with *RAP1* dosage and environment specific conditions

To further explore gene expression patterns associated with *RAP1* dosage across different environments, we applied Weighted Gene Co-expression Network Analysis (WGCNA)^41^. This approach clusters co-expressed genes into modules based on their expression profiles and quantifies the correlation between these modules and environmental conditions. Our analysis identified four distinct co-expressed gene modules named Module 1 – Module 4 (Figure 5A). 1025 genes which were not assigned to any of these four modules were grouped into Module 0.

**Figure 5.**
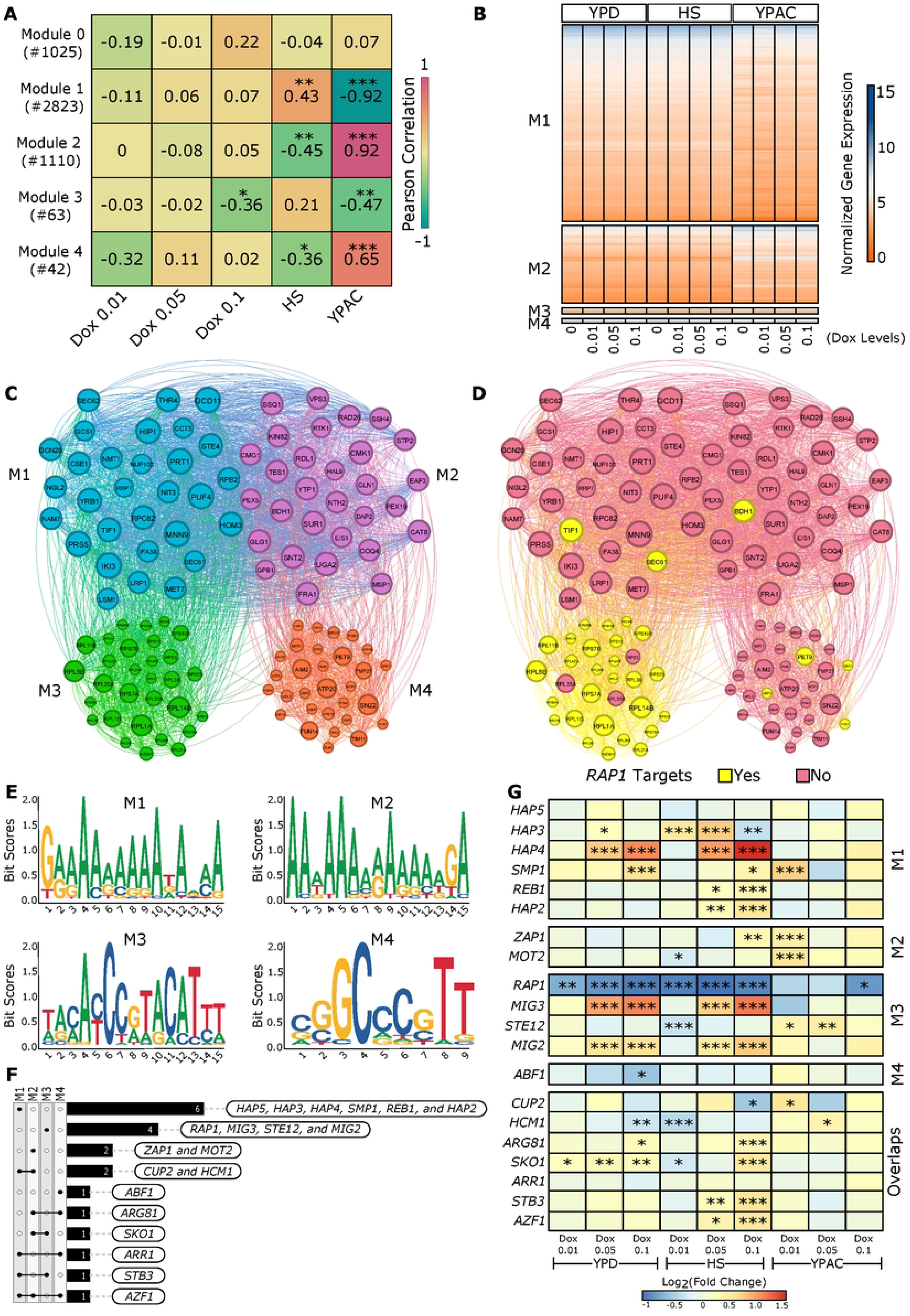
Network modules and regulatory signatures. A) Heatmap showing Pearson correlations between module membership and sample metadata. Numbers in parentheses indicate the number of genes in each module. B) Heatmap of normalized expression values for genes in each module across different *RAP1* levels and environments. C) Network view of the top 30 hub genes from each module. D) Same network annotated with known *RAP1* downstream targets. E) MEME analysis highlighting promoter nucleotide enrichment for genes in the four modules. F) Overlap of transcription factor enrichments identified across modules. G) Heatmap of log2 fold change values for enriched transcription factors. Asterisks denote significant changes relative to maximal *RAP1* level in that environment.

We performed Pearson correlation analysis between module eigengenes (the first principal component of each module’s expression matrix, summarizing the overall expression trend within that module) and various experimental conditions such as *RAP1* dosage and environmental context (YPD, HS, and YPAC) (Figure 5A). Module 0 genes showed no statistically significant correlation with any condition; therefore, genes in Module 0 were excluded from downstream analysis. Module 1, comprising 2823 co-expressed genes, showed a significant positive correlation with the HS environment and a strong negative correlation with YPAC. This highlights that genes in Module 1 tend to be upregulated under heat shock and downregulated in the acetate-rich YPAC environment (Figure 5A, B). In contrast, Module 2, which includes 1110 genes, exhibited the opposite trend, a significant negative correlation with HS and a strong positive correlation with YPAC (Figure 5A, B). These reciprocal patterns between Module 1 and Module 2 highlight environment-specific gene expression programs that respond oppositely in HS and YPAC. Module 3, consisting of 63 co-expressed genes, showed a significant negative correlation with both Dox 0.1 and the YPAC environment, indicating that genes in this module are downregulated in response to the lowest *RAP1* levels (Dox 0.1) tested and in acetate conditions (Figure 5A, B). Module 4, which contains 42 genes, also showed a significant negative correlation with HS and a moderate positive correlation with YPAC, further underscoring transcriptional shifts associated with environmental changes (Figure 5A, B).

### *RAP1* Targets Are Enriched Among Highly Connected Genes in Module 3

Next, we aimed to identify the most highly connected genes within each module (referred to as hub genes) to visualize the underlying network structure. For each module, we selected the top 30 hub genes based on their absolute module membership (kME) values (see methods), which reflect how strongly each gene is associated with its respective module. The resulting network plot represents genes as nodes, with node size proportional to their degree (the number of connections) and node color corresponding to their module assignment (Figure 5C). To explore potential regulatory relationships, we overlaid a curated list of known *RAP1* target genes onto these hub gene networks. Notably, we found that module 3 contained the highest number of *RAP1* targets (Figure 5D), suggesting that *RAP1* may play a central role in directly regulating the genes within this module.

To investigate potential transcriptional regulators associated with each gene co-expression module, we analyzed the promoter regions (−500 bp upstream) of the top 30 hub genes from each module for enriched DNA sequence motifs. This analysis was performed separately for each module to capture module-specific regulatory patterns. Using YEASTRACT^46^ as a reference database, we identified 20 potential transcription factors (TFs) associated with the enriched motifs, thereby linking each module to candidate upstream regulators. Modules 1 and 2 were enriched for A/T-rich motifs while Module 3 displayed a complex enriched motif indicating a more specific TF binding pattern (Figure 5E). Module 4 was enriched for a distinct G/C-rich motif pattern (Figure 5E).

Predicted TFs revealed module specific as well as shared regulators. TFs such as *HAP5*, *HAP3*, *HAP4*, *SMP1*, *REB1*, and *HAP2* were exclusively predicted for module 1 (Figure 5F). *ZAP1* and *MOT2* were unique to module 2 (Figure 5F). TFs including *RAP1*, *MIG3*, *STE12*, and *MIG2* were exclusive to module 3. *RAP1* enrichment in module 3 was expected as our network analysis pointed out that most of the module 3 hub genes were direct *RAP1* downstream targets (Figure 5F). For module 4, *ABF1* was predicted as a key regulator. Several TFs were shared across modules, pointing to broader regulatory roles: *CUP2* and *HCM1* were common to modules 1 and 2. *SKO1* to modules 2 and 3. *STB3* to modules 1 and 3. *ARR1* to modules 1 and 4 and *ARG81* to Modules 2 and 4. Finally, TF *AZF1* was predicted to regulate genes across modules 1, 2, and 4 (Figure 5F). Because the predicted transcription factors for these modules likely contribute to regulating the observed gene-expression changes, we examined how their own expression varied across *RAP1* levels and environmental contexts. To do that, we looked at the log_2_ fold-change of all 20 predicted TFs in our RNA-seq data and found that 18 out of these 20 exhibited significant expression changes in at least one of the tested conditions (Figure 5G).

## Discussion

In this study, using the Tet-Ou system, we generated distinct *RAP1* expression levels across three conditions: fermentative growth in YPD, HS, and respiratory growth in YPAC (Figure 1). Our previous work identified a small set of Dox-responsive genes (∼30)^47^; these were removed from the present analysis to focus exclusively on the euects of *RAP1* depletion. Even at baseline (maximal *RAP1* expression, no Dox), the three environments produced markedly diuerent transcriptional landscapes (Figure 2A-F). YPAC showed extensive transcriptional remodeling relative to YPD, with 3727 DEGs, which were primarily associated with respiration, mitochondrial biogenesis, and proton transmembrane transport while heat shock produced a smaller response comprising 806 DEGs (S-Table 4). Most heat shock associated genes were upregulated, and their GO enrichment terms were consistent with activation of stress-induced chaperones, iron transport, protein-folding pathways, and other canonical heat response mechanisms (S-Table 4). The YPAC specific DEGs showed strong enrichment for genes previously reported to respond to acetate (enrichment score 0.67), indicating close agreement with established literature. Similarly, the HS gene set showed positive enrichment with established heat-responsive genes (score 0.40), confirming that our experimental conditions elicited expected stress response modules (Figure 2 H, I). We also identified a set of 564 genes whose expression was sensitive to changes in environment and identified in every environmental comparison, pointing to a core group of loci that integrate metabolic and stress cues (Figure 2G).

When we investigated the known regulatory functions of *RAP1* across environments, we found a significant change in gene expression of direct *RAP1* targets. The mean expression values (in FPKM) of direct *RAP1* target genes were 1450.50 in YPD, 1449.19 in HS, and 987.79 in YPAC, suggesting a significant reduction in gene expression in YPAC compared to both HS and YPD (Figure 2J). This trend is further reinforced by the coordinated downregulation of RPGs in YPAC with respect to both HS and YPD, which are known to be tightly linked to cellular growth rate (mean FPKM YPD: 3257.88, HS: 3330.2, and YPAC: 2071.67) (Figure 2K). Telomeric genes also exhibited lower expression in YPAC (mean FPKM: 15.09) compared to YPD (32.35) and HS (25.87), consistent with the pattern observed for *RAP1* targets and RPGs (Figure 2L). In contrast, glycolytic genes (mean FPKM: YPD 1958.08, HS 1759.31, YPAC 1956.84), DNA repair genes (YPD 517.86, HS 513.94, YPAC 377.94), and SML genes (YPD 137.94, HS 148.50, YPAC 251.11) did not show significant changes in expression across the three conditions (Figure 2M-O). This overall suggests that *RAP1’s* regulatory role is highly context-dependent, and the metabolic and/or stress state of the cell shapes how *RAP1’s* regulatory network is structured and deployed.

In both YPD and HS conditions, progressive depletion of *RAP1* led to an increasing number of DEGs, indicating a dose-dependent transcriptional response (Figure 3A-L). In contrast, the YPAC condition exhibited a distinct and inverse pattern, where decreasing *RAP1* levels resulted in fewer DEGs (Figure 3M-R, 4E). This suggests that the transcriptional network’s sensitivity to *RAP1* is fundamentally altered under non-preferred carbon sources like acetate, likely due to metabolic reprogramming. Our analysis reveals extensive condition-dependent transcriptional reprogramming across growth and stress environments. Most genes did not maintain a consistent regulatory direction across YPD, HS, and YPAC, indicating that gene regulation is highly plastic rather than fixed (Figure 4F). Notably, a substantial fraction of genes (1244) was upregulated under multiple conditions (Figure 4F). In contrast, relatively few genes were uniformly downregulated (289). Genes that switch regulatory direction between conditions likely reflect metabolic and physiological trade-ous, where growth/survival associated programs are suppressed in rich media and activated under stress. Together, these results support a model in which environmental cues drive targeted transcriptional rewiring, enabling cells to balance growth and survival through flexible regulatory control (Figure 4F) highlighting the plasticity of transcriptional regulation of yeast where both genetic (*RAP1* levels) and environment (carbon source, temperature) interact to shape gene expression.

WCGNA further elucidated the structure of GXE interactions by identifying modules of co-expressed genes with distinct environments and *RAP1* level associations (Figure 5A). Overall, Module 1 and Module 2 showed strong, environment-specific expression patterns, but in opposite directions between HS and YPAC. Module 3 is moderately downregulated by the lowest *RAP1* dosage (Dox 0.1) and in YPAC. Module 4 is similar to Module 2’s expression pattern, up in YPAC, down in HS (Figure 5A).

Module 1 is primarily associated with ribosome biogenesis, rRNA/tRNA processing, and related biosynthetic processes. Its enriched molecular functions include catalytic activity on RNA and tRNA, RNA polymerase activity, and transmembrane transporter functions (S-Table 6). Enrichment in ribosomal and endoplasmic reticulum components further supports a strong link to ribosome assembly and RNA processing, highlighting Module 1’s core role in the protein synthesis machinery. Module 2 is dominated by energy metabolism, including cellular respiration, oxidative phosphorylation, and various catabolic processes. Its enriched molecular functions highlight electron transfer activity, oxidoreductase functions, and active transmembrane transport. The cellular component enrichments are centered on microbodies, peroxisomal membranes, and mitochondrial structures (S-Table 6), indicating that Module 2 plays a key role in cellular energy production through mitochondrial and peroxisomal metabolic pathways. Module 3 is focused on rRNA metabolism, ribosome assembly, translation, and nuclear export as its primary biological processes. Enriched molecular functions include structural molecule activity, particularly related to ribosomal components. Cellular component enrichment highlights ribosomal subunits and associated structures (S-Table 6), indicating that Module 3 extends the theme of ribosome biogenesis with a specific emphasis on ribosome assembly and export processes critical for protein synthesis. Module 4 is enriched for biological processes related to nucleotide and ribonucleotide metabolism and biosynthesis. Its molecular functions include proton and cation channel activity, passive transmembrane transport, and ligase activity. Cellular component enrichment highlights mitochondrial membranes, respiratory chain complexes, and transporter complexes (S-Table 6). Together, these features suggest that Module 4 is specialized for mitochondrial function and nucleotide metabolism, particularly supporting ATP synthesis and membrane transport.

The identification of candidate TF(s) regulating these modules, and the analysis of their expression patterns provide a potential mechanistic basis for the observed GXE interactions. TFs predicted for module 1, including *HAP2, HAP3, HAP4, and HAP5*, function as global regulators of respiratory gene expression (Figure 5F, G). *SMP1* and *REB1* are associated with the cellular response to osmotic stress^48–50^ and with termination of transcription by RNA polymerase I^51,52^, respectively. In module 2, *ZAP1* regulates transcription^53^, whereas *MOT2* acts as a global regulator by modulating mRNA turnover and protein ubiquitination under stress conditions^54^. *MIG2* and *MIG3*, also assigned to this module, are transcriptional repressors of glucose-responsive genes^55,56^. Consistent with this role, these genes are downregulated under acetate conditions compared to heat stress, promoting cellular adaptation and survival (Figure 5F, G). *STE12*, a regulator of invasive growth^57^ in response to glucose limitation, is upregulated in acetate relative to heat stress. Finally, *ABF1*, predicted for module 4, is implicated in chromatin reorganization and participates in transcriptional activation, gene silencing, and DNA replication and repair^58^ (Figure 5F, G).

While our study provides a comprehensive analysis of *RAP1* dependent transcriptional responses across three environments, it does not capture post-transcriptional or translational regulatory layers that may further modulate phenotypic outcomes. Our findings reinforce the importance of studying gene regulation across multiple environmental contexts to fully elucidate the architecture of GXE interactions. Traditional approaches that examine gene expression in a single environment may overlook critical context specific regulatory mechanisms. Use of high throughput transcriptomics and network analysis provides a systems-level perspective on how transcriptional networks reconfigure in response to combined genetic and environmental perturbations. This approach can be extended to other TFs and environmental variables, ouering a blueprint for dissecting the complexity of regulatory networks in diverse organisms.

## Materials and Method

### Sample generation

The yeast strains used in this study were obtained from the Yeast Tet Promoter Hughes Collection (yTHC), obtained from Horizon/Dharmacon Reagents^40^. Specifically, we used the TetO7-*RAP1* strain^35^, in which the endogenous *RAP1* promoter was replaced with a Tet-titrable promoter (Figure 1A). This allows for the regulation of *RAP1* expression by the presence and abundance of Dox in the growth media (Figure 1B). This TetO_7_-*RAP1* strain was first grown overnight in YPD medium to saturation for approximately 24 hours (Figure 1C). The overnight cultures were then diluted with fresh YPD liquid medium and grown to mid-exponential phase for 5–8 hours. At this stage, the cultures were further diluted in fresh YPD medium containing a range of Dox concentrations (0, 0.01, 0.05, and 0.1 µg/mL) and incubated at 30°C with constant shaking at 200 RPM for 12–14 hours, or until the cultures reached an optical density (OD) of 0.6–0.8 (Figure 1C). OD measurements were performed using a BioPhotometer Plus at a wavelength of 600 nm. The cultures were then pelleted by centrifugation, the excess media was removed, and the cell pellets were snap-frozen in liquid nitrogen and immediately stored at −80°C. For HS treatment, the TetO_7_-*RAP1* strain was grown following the same protocol as described for YPD cultures. However, after reaching mid-exponential phase, the cultures were subjected to heat shock by incubation at 37°C for 15 minutes. For YPAC samples, the YPD medium was replaced with YPAC medium, and the same growth and collection steps were followed (Figure 1C).

### RNA extraction and sequencing

The cell culture pellets were incubated in a 100 μL mixture containing 50 U of 20 T lyticase at 30 °C for 30 min. After incubation, total RNA was extracted using the SV Total RNA Isolation System (Promega) with a modified protocol^59^. The recovered RNA was treated for 15 min with DNase at room temperature. Samples were sent to the University of Michigan Sequencing Core where barcoded cDNA libraries were created and sequenced on a lane of an Illumina HiSeq 4000 for 76 cycles using single end sequencing. Three replicates per sample were sequenced for three environments and four dox levels resulting in 36 sequencing files (fastq). The sequencing depth and mapping percentages of these fastq files are mentioned in S-Table 7. The sequencing data was then analyzed using the RNA sequencing pipeline outlined in Figure 1D.

### BIOL310: Genomics analysis performed in a Course-based Undergraduate Research Experience (CURE)

The downstream bioinformatics analysis for this RNA-seq project was carried out by 22 undergraduate students enrolled in BIOL310: Genomics Analysis, a semester-long course at Wesleyan University. This manuscript is the sixth generated through this course^35,60–63^ over the past six years, which is designed to provide early-stage undergraduates with authentic, hands-on research experience and direct engagement in scientific discovery. Students work with previously unanalyzed RNA-seq datasets and apply contemporary genomics and bioinformatics tools in a discovery-driven framework. All students contributed to every stage of the project, including quality control, bioinformatics processing, statistical analysis, and interpretation of results. The manuscript text reflects the integration of their individual analyses, ideas, and interpretations into a single, cohesive study.

As part of this CURE-style course, students conducted analyses using the Galaxy platform (https://usegalaxy.org/), followed by downstream data visualization. The complete analysis pipeline is shown in Figure 1D. Raw FastQ files generated by the University of Michigan Sequencing Core Facility were first subjected to quality control using FastQC^64^. Reads were then aligned to the *Saccharomyces cerevisiae* reference genome (Saccharomyces_cerevisiae.R64–1-1.dna.toplevel.fa), obtained from the Ensembl genome browser, using Bowtie2^65^ with default parameters, producing BAM files. Gene expression was quantified with Cufflinks^66^ using the corresponding GFF3 annotation file (Saccharomyces_cerevisiae.R64–1-1.105.gff3). Expression values were normalized for gene length and sequencing depth and reported as fragments per kilobase per million reads (FPKM). Differential gene expression was assessed using Cuffdiff^66^, and results were visualized and analyzed in R using RStudio.

### Gene Set Enrichment analysis

Gene sets used for enrichment comparisons were obtained from AllianceMine, which is linked to the Saccharomyces Genome Database (SGD)^67^. For each condition, all genes annotated to the specified GO term, as well as to its child terms, were retrieved. The GO query “heat” was used to define the heat shock gene set, while the query “acetate” was used to identify genes associated with acetate-related conditions. The data was further filtered down to keep genes belonging to *S. cerevisiae* (S-Table 3). Gene set enrichment analysis was performed in R using the fgsea package (v1.32)^68^.

### Gene Ontology analysis

To functionally characterize genes, we performed GO enrichment analysis using the clusterProfiler^69^ R package (v4.14.4). Library(org.Sc.sgd.db) was used as an annotation database for *S. cerevisiae*. Each gene set was analyzed for Biological Process (BP), Molecular Function (MF), and Cellular Component (CC). Enrichment was assessed using a hypergeometric test with Benjamini-Hochberg (BH) correction^70^ for multiple testing, applying a p-value cutoff of 0.05.

### Principal Component Analysis (PCA)

To evaluate overall transcriptional variation across conditions, PCA was performed using the PCAtools^71^ (v2.18.0) R package. Gene expression matrices were generated by merging Cufflinks-processed data across all *S. cerevisiae* samples under YPD, HS, and YPAC environments with varying Dox concentrations. All expression values were log_2_ transformed (log_2_(FPKM+0.001)). PCA was conducted by excluding the bottom 10% of low variance genes.

### Weighted Gene Co-expression Network Analysis (WGCNA) and Network visualization

WGCNA analysis was performed in R using the WGCNA package^41^. Cufflinks output files from all 36 samples were merged to generate a single input expression matrix. FPKM values were log_2_ transformed using the formula log_2_(FPKM + 1) to normalize the data. Sample and gene quality check were conducted using the goodSamplesGenes() function, which confirmed all samples passed the quality filter. A total of 67 low-quality genes were flagged and subsequently removed, resulting in a final dataset of 5063 genes. We constructed a signed network using a soft-thresholding power (β) of 20, selected based on the scale-free topology criterion (R² > 0.80).

Co-expression network was constructed using the blockwiseModules() function with the following parameters: TOMType = “signed”, power = 20, mergeCutHeight = 0.25, and maxBlockSize = 6000. Modules were detected using hierarchical clustering of the topological overlap matrix (TOM) and dynamic tree cutting. Modules were assigned numeric labels, and the module eigengenes (MEs) were extracted to represent each module (Figure 5A). To assess associations between modules and experimental metadata, we binarized categorical variables representing environmental condition (YPD, HS, and YPAC) and Dox treatments (Dox 0, 0.01, 0.05, and 0.1 µg/mL) using the binarizeCategoricalColumns() function. Pearson correlations between module eigengenes and these binarized traits were calculated using the base cor() function, and associated p-values were computed using corPvalueStudent() with the number of samples as input. Module-trait relationships were further visualized using the function CorLevelPlot().

For network visualization, we identified highly connected intramodular hub genes by calculating the module membership (kME) values. kMEs are the individual gene expression correlation to the corresponding MEs, for all genes within each module. Top 30 genes ranked by absolute kME were selected, resulting in a final set of representative hub genes across all detected modules (Figure 5C-D).

### Motif enrichment and Transcription Factor (TF) prediction

Promoter sequences for the top 30 hub genes in each module were extracted using a custom in-house R script. For each module, the resulting FASTA files containing promoter sequences were submitted to the XSTREME tool^72^ (part of the MEME Suite^73^) for de novo motif discovery and enrichment analysis. The YEASTRACT database^46^ was used as the reference. From each module, the top-ranked enriched motif identified by XSTREME was selected for visualization, and the predicted TF list (using TOMTOM^74^) was complied with respect to all motifs that were found enriched for each module.

## Data availability

The sequence data generated in this work is available at the NCBI Gene Expression Ominibus (GEO) under accession (available at the time of publication) and the code for downstream analysis will be available at github (at the time of publication).

## Authorship contribution statement

**SK:** Conceptualization, Methodology, Software, Formal analysis, Investigation, Data curation, Writing – original draft, Writing – review & editing, Visualization, Supervision. **GS:** Conceptualization, Methodology, Software, Formal analysis, Investigation, Data curation, Writing – original draft. **AS:** Formal analysis, Investigation, Data curation, Writing – original draft, Visualization for YPD environment. **AB:** Formal analysis, Investigation, Data curation, Writing – original draft, Visualization for YPD environment. **BO:** Formal analysis, Investigation, Data curation, Writing – original draft, Visualization for YPD environment. **IB:** Formal analysis, Investigation, Data curation, Writing – original draft, Visualization for YPD environment. **MM:** Formal analysis, Investigation, Data curation, Writing – original draft, Visualization for YPD environment. **SC:** Formal analysis, Investigation, Data curation, Writing – original draft, Visualization for YPD environment. **SP:** Formal analysis, Investigation, Data curation, Writing – original draft, Visualization for YPD environment. **AL:** Formal analysis, Investigation, Data curation, Writing – original draft, Visualization for HS environment. **YZ:** Formal analysis, Investigation, Data curation, Writing – original draft, Visualization for HS environment. **GB:** Formal analysis, Investigation, Data curation, Writing – original draft, Visualization for HS environment. **LP:** Formal analysis, Investigation, Data curation, Writing – original draft, Visualization for HS environment. **MS:** Formal analysis, Investigation, Data curation, Writing – original draft, Visualization for HS environment. **MN:** Formal analysis, Investigation, Data curation, Writing – original draft, Visualization for HS environment. **SP:** Formal analysis, Investigation, Data curation, Writing – original draft, Visualization for HS environment. **BM:** Formal analysis, Investigation, Data curation, Writing – original draft, Visualization for YPAC environment. **CP:** Formal analysis, Investigation, Data curation, Writing – original draft, Visualization for YPAC environment. **EP:** Formal analysis, Investigation, Data curation, Writing – original draft, Visualization for YPAC environment. **EJ:** Formal analysis, Investigation, Data curation, Writing – original draft, Visualization for YPAC environment. **EH:** Formal analysis, Investigation, Data curation, Writing – original draft, Visualization for YPAC environment. **PH:** Formal analysis, Investigation, Data curation, Writing – original draft, Visualization for YPAC environment. **TT:** Formal analysis, Investigation, Data curation, Writing – original draft, Visualization for YPAC environment. **JC:** Conceptualization, Methodology, Software, Resources, Writing – original draft, Writing – review & editing, Supervision, Project administration, Funding acquisition.

## Acknowledgements

Research reported in this publication was supported by Wesleyan University (start-up funds to J.D.C. and Department of Biology funds to J.D.C.), the Wesleyan Ronald E. McNair Post Baccalaureate Program (G.S.) and the National Institute of General Medical Sciences of the National Institutes of Health under Award Number R15GM135901 (awarded to J.D.C.). The content is solely the responsibility of the authors and does not necessarily represent the official views of the National Institutes of Health.

## Supplementary

**S-Figure1:** Overlap of shared GO enriched terms in different comparisons (HS vs YPD, YPAC vs YPD, and YPAC vs HS) at maximum *RAP1* levels.

**S-Figure2:** Overlap of shared GO enriched terms at all *RAP1* levels in YPD, HS and YPAC environment.

## Table Legends

### Supplementary

**S-Table1:** Gene expression (FPKM) values across various *RAP1* levels in YPD, HS and YPAC environment.

**S-Table2:** List of overlapping significant DEGs at max *RAP1* level

**S-Table3:** List of Heat and Acetate genes downloaded from SGD used for GSEA

**S-Table4:** Gene ontology enrichment terms observed at max *RAP1* levels in YPD, HS and YPAC

**S-Table5:** Gene ontology enrichment terms observed at various *RAP1* levels in YPD, HS and YPAC

**S-Table6:** Gene ontology enrichment terms for genes belonging to different network modules.

**S-Table7:** Summary of sample collection, RNA sequencing depth and mapping percentages.

